# Severity of Early Life Stress Moderates the Effect of Fine Particle Air Pollution on Adolescent Brain Development

**DOI:** 10.1101/763896

**Authors:** Jonas G. Miller, Emily L. Dennis, Booil Jo, Ian H. Gotlib

## Abstract

Air pollution is currently the greatest environmental threat to public health, but we know little about its effects on adolescent brain development. In this context, exposure to air pollution co-occurs, and could interact, with social factors that also affect brain development, such as early life stress (ELS). Here, we show that severity of ELS moderates the association between fine particle air pollution (particulate matter 2.5; PM2.5) and structural brain development. We interviewed adolescents about ELS, used census-tract data to characterize PM2.5 concentrations, and conducted longitudinal tensor-based morphometry to assess regional changes in brain volume over a two-year period. Across various cortical, thalamic, and white matter tract regions, there was a remarkably consistent effect of PM2.5 on volumetric change for adolescents who had experienced less, rather than more, severe ELS. Furthermore, exposure to higher levels of PM2.5 and experiencing moderate to severe ELS were associated with comparable volumetric changes in the brain in adolescence.

---

Air pollution is currently the greatest environmental threat to public health (1). Exposure to high levels of air pollution is associated with risk for respiratory infection, asthma, lung cancer, heart disease, stroke incidence, and Alzheimer’s and Parkinson’s disease (2, 3); moreover, air pollution reduces average global life expectancy by approximately two years per person (4). Particulate matter 2.5 (PM2.5), or air particles smaller than 2.5 µm in diameter, may be especially dangerous. PM2.5 particles can be composed of harmful mixtures of metals and chemicals that are easily inhaled deeply into the lungs and absorbed into the bloodstream, leading to local and systemic inflammation and oxidative stress (2) which, in turn, can adversely affect brain development (5). Smaller PM2.5 particles can also reach the brain, potentially contributing to neuroinflammation and brain damage. Ultrafine particles originating from combustion processes can enter the brain directly via the olfactory nerve and bulb (6), as well as indirectly after entering the blood stream and translocating across the blood-brain barrier (7). Despite these findings, compared to our understanding of how PM2.5 affects cardiorespiratory functioning and physical health, we still know far less about the impact of PM2.5 on pediatric brain development in humans, especially beyond early childhood (5).

Adolescents may be particularly vulnerable to the effects of PM2.5 on brain development. Compared to adults, adolescents spend more time being physically active outdoors, increasing their exposure to ambient PM2.5 (8). Adolescence is also a period of rapid and dramatic brain development (9), which may confer greater vulnerability to environmental insults. A growing number of studies are reporting evidence that PM2.5 exposure has negative effects on cognitive and affective functioning in children and adolescents (10–13). Fewer studies, however, have investigated the impact of PM2.5 on the adolescent human brain. Much of the existing neuroimaging research on air pollution has used cohorts from the Mexico City Metropolitan Area. For example, increased exposure to PM2.5 in youth from Mexico City has been associated with damage to the nasal epithelium and blood-brain barrier breakdown, potentially allowing more PM2.5 particles as well peripheral proinflammatory cytokines to pass into the brain (14). These youth also show white matter damage associated with increased neuroinflammation (15), further suggesting that PM2.5 exposure has adverse effects on child and adolescent brain health. Other research has also found links between PM2.5 exposure and altered brain structure and function. Prenatal exposure to higher levels of air pollution has been associated with greater cortical thinning and reduced white matter volume, which, in turn, were associated with cognitive impairment (12, 16). In a cross-sectional study of 8- to 12-year-old children in Spain, increased exposure to traffic-related air pollution was associated with less functional integration and segregation of the default mode brain network (17).

Collectively, these findings suggest that exposure to PM2.5 adversely affects adolescent brain structures and networks that are critical for healthy cognitive and affective functioning. It is important to note, however, that previous studies either have considered the brain only at a single time point or have assessed the effects of PM2.5 on change in brain structures in youth over one year. It is unclear how PM2.5 may affect adolescent brain development over a longer interval. In addition, studies of adolescents have primarily used cohorts from heavily air polluted metropolitan areas. For example, half of people living in the Mexico City metropolitan area are exposed to at least 24 µg/m^3^ mean daily PM2.5 (18); in contrast, many metropolitan areas in the United States are below the national standard for annual average of 12 µg/m^3^ set by the Environmental Protection Agency (EPA). We know less about whether PM2.5-related effects on adolescent brain development are detectable in youth living in these less air polluted areas.

Although PM2.5 may alter brain structure and function directly, it is important to recognize that exposure to air pollution co-occurs in the context of different social environments and experiences that are also implicated in adolescent brain development. Consistent with this perspective, some researchers have advocated for examining the joint contributions of environmental and psychosocial factors to health (19–22). Assessing the interaction of early life stress (ELS) and PM2.5 may be particularly promising for gaining a more comprehensive understanding of variability in adolescent brain development that may be linked to health. ELS has consistently been shown to have deleterious effects on the brain (23) and to contribute to pathophysiology, such as proneness to exaggerated proinflammatory responses (24), that could modify the impact of PM2.5 on risk for disease (22). Indeed, findings of studies of child and adolescent respiratory health indicate that susceptibility to the adverse effects of air pollution may depend on exposure to ELS. Specifically, there is evidence that ELS increases susceptibility to the adverse effects of traffic-related air pollution on the development of asthma and lung functioning (25–27). Taken together, these findings suggest a “double exposure” model in which ELS exacerbates risk related to increased air pollution. Alternatively, it is possible that youth exposed to less severe ELS will be more sensitive to the effects of air pollution. For example, particularly high levels of stress or air pollution could exacerbate risk regardless of the level of the other factor (20). In fact, in a study of asthmatic children living in urban public housing, improving air quality by reducing allergen exposure was less effective in reducing symptoms for families who reported greater fear of violence in their communities (28). This finding fits with a saturation effect – that is, stress may overpower positive or negative contributions related to air quality (20). From a somewhat different perspective, ELS may organize or calibrate neurobiology in a way that reduces the impact of subsequent stressors, including those from the physical environment (29, 30). More specifically, both theoretical and empirical work suggests that children exposed to moderate and severe ELS are less sensitive to social environmental influences on behavioral functioning than are their peers exposed to minimal ELS (29); however, this research has not considered whether ELS moderates sensitivity to air pollution exposure.

The present study was conducted to address the question of whether there are interactive effects between ELS and PM2.5 that conform to a double exposure, saturation effect, or calibration model. Specifically, we examined whether individual differences in severity of ELS moderate the association between neighborhood-level PM2.5 and adolescent brain development. In a sample of adolescents living in the San Francisco Bay Area, we used longitudinal tensor-based morphometry (TBM) to assess expansion and contraction patterns across the brain that were associated with the interaction of PM2.5 and ELS severity. Given that no previous study has considered how PM2.5 and ELS jointly affect brain development, we did not generate *a priori* hypotheses about, or constrain analyses to, specific brain regions. Rather, using longitudinal TBM allowed us to explore which brain structures were associated with the interaction of PM2.5 and ELS. Other types of FreeSurfer-based cortical analyses that focus on surface area and cortical thickness (31) do not allow for whole-brain (including white-matter) voxel-wise analysis of brain volume. TBM has several advantages over voxel-based morphometry (VBM) (32), which is also a whole-brain analytic approach. Unlike VBM, TBM does not involve tissue segmentation or spatial smoothing steps, which can introduce error and limit resolution for small effects. In addition, VBM depends on tissue density measurements to calculate volume differences, which require high-quality registration. Thus, inaccuracies in registration can mistakenly be identified as volume differences (33).

We evaluated three different hypotheses regarding the nature of interaction effects. The double exposure model would predict that PM2.5 will be associated with patterns of brain development most strongly in adolescents who experienced more severe ELS. In contrast, the saturation effect model would predict that more severe experiences of ELS will overpower any association with PM2.5. Thus, potentially problematic patterns of brain development related to higher levels of PM2.5 will be observed for adolescents who experienced less severe ELS, and ELS will have a robust association with volumetric changes. Finally, a calibration perspective would predict that youth exposed to more severe ELS will be less sensitive to the negative effects of PM2.5 on brain development than will their peers exposed to milder ELS.

## Results

As part of a larger study on early life stress and neurodevelopment, 116 adolescents (56% girls) from the San Francisco and San Jose Bay Area completed an interview to assess ELS severity and provided useable TBM data at Time 1 (Mean age=11.50; SD=1.08) and two years later at Time 2 (Mean age=13.47; SD=1.16) of the study. Participants were diverse in terms of race and ethnicity (51% White; 11% Asian; 8% Hispanic/Latino; 5% Black; 19% More than one race or ethnicity; 4% Other; 2% Did not report) and family income (58% of families reported an annual income over $100K; range from less than $5k to greater than $150K). We used air quality data compiled and made publicly available by the CalEPA to assess PM2.5 concentrations for each participant’s census tract. The majority of participants lived in unique census tracts (i.e., 116 participants in 106 unique census tracts); thus, we expect little effect of clustered participants in the same neighborhood. Descriptive statistics and correlations are presented in the supporting information in Table 1. Of the 214 adolescents who participated in the larger study, those who did not consent to be scanned at either time point (n=35), did not provide useable TBM data due to motion (based on visual inspection) (n=31), had braces (n=16), dropped out of the study after T1 (n=15), or did not have PM2.5 data (n=1), were not included in our analyses.

**Table 1.**
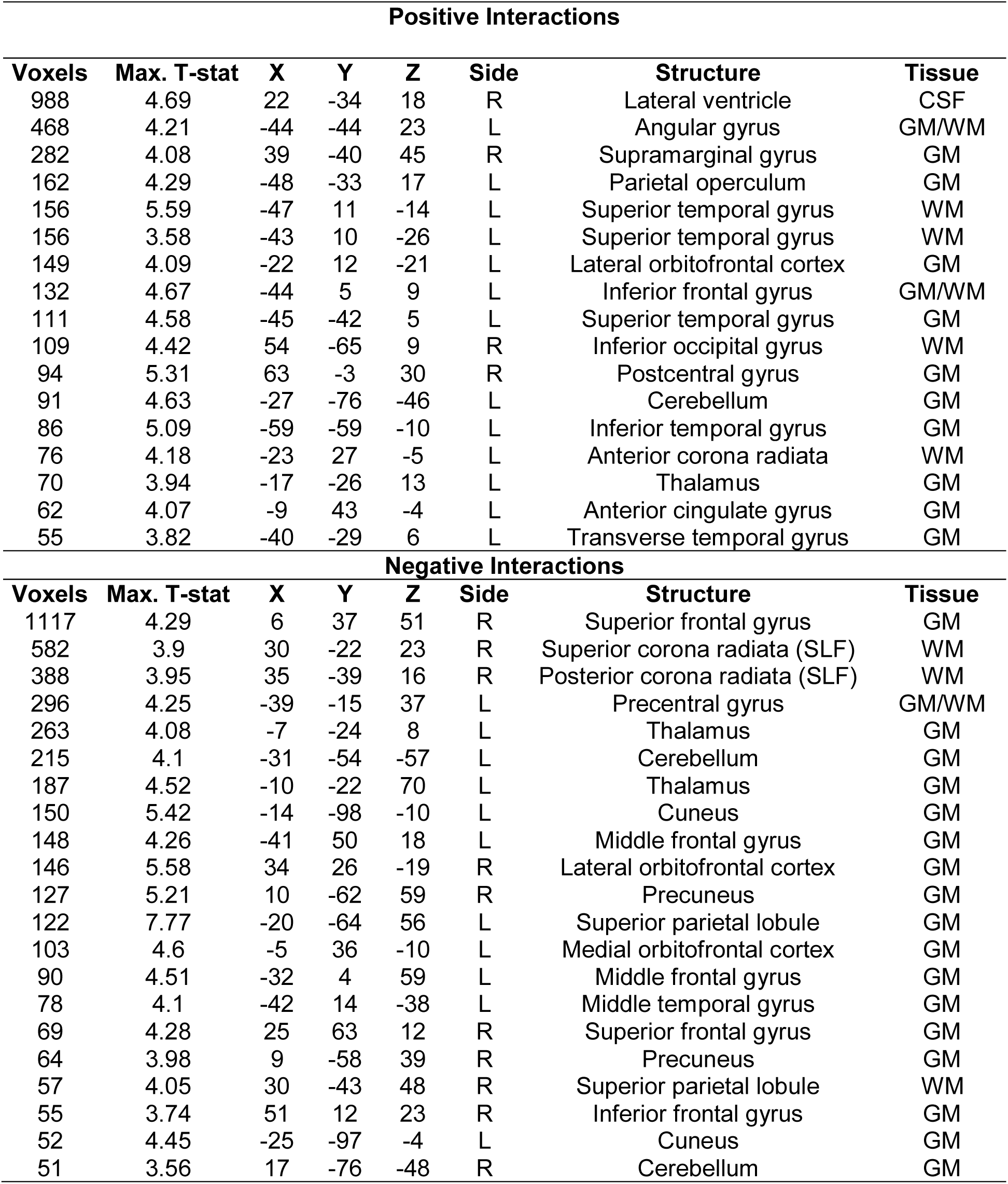
Clusters showing a significant interaction between ELS and PM on regional brain volume. Note. Shown are the cluster size, peak regression statistic, coordinates (MNI), hemisphere, structure, and tissue type. R=right; L=left; GM=grey matter; WM=white matter; CSF=cerebrospinal fluid.

Our longitudinal whole-brain analysis covaried for sex, age at Time 1, interval between Time and Time 2, total brain volume at Time 1, and neighborhood socioeconomic disadvantage. Sensitivity analyses showed that there were minimal difference between models in which we excluded versus included different covariates (e.g., minority status) and indexed SES using separate multiple covariates (see supplementary materials for details). There were no significant main effects of ELS severity or PM2.5 on regional changes in brain volume after adjusting for multiple comparisons. Conversely, there were significant interactions between ELS and PM2.5 on 38 clusters larger than 50 voxels (see Figure 1 and Table 1). From these clusters we extracted change estimates averaged within each cluster. After inspecting and removing extreme outlier TBM values for each cluster (see supplementary materials for details), 26 clusters remained. There were positive associations between the ELS by PM2.5 interaction term and change in brain volume in the lateral ventricle, lateral orbitofrontal cortex, anterior corona radiata white matter tract, and a number of predominantly grey matter clusters across the frontal temporal, parietal, and occipital cortices (14 total; see Figure 2). There were negative associations between the interaction term and change in brain volume in the superior corona radiata white matter tract, thalamus, and a number of predominantly grey matter clusters across the frontal, temporal, parietal, and occipital cortices (12 total; see Figure 3).

**Figure 1.**
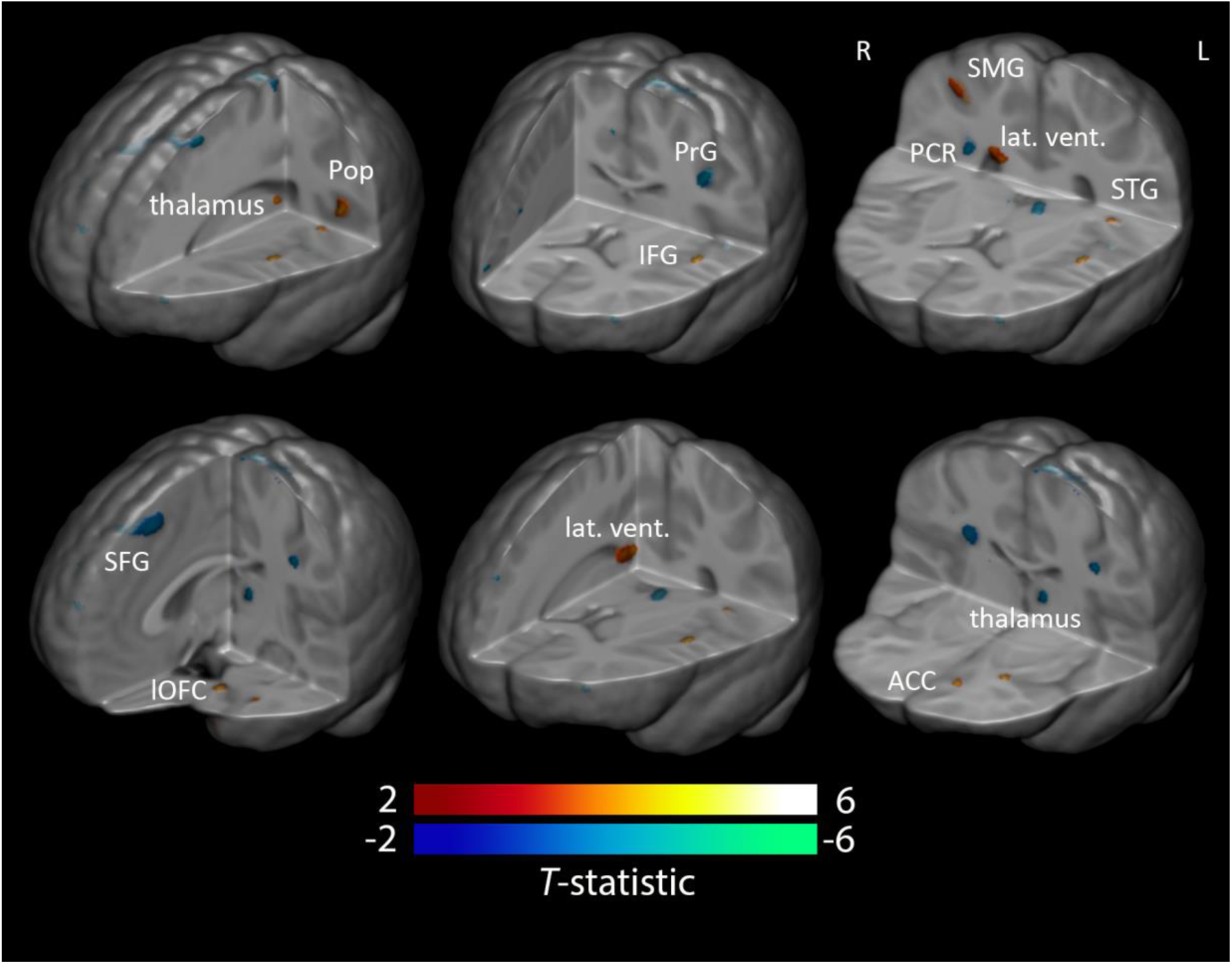
Clusters showing a significant interaction between ELS and PM on regional brain volume. Note. T-maps are shown for significant clusters, with blue-green for negative associations and red-yellow for positive associations. Left in image is right in brain. Pop=parietal operculum, PrG=precentral gyrus, IFG=inferior frontal gyrus, PCR=posterior corona radiata, SMG=supramarginal gyrus, lat. vent.=lateral ventricle, STG=superior temporal gyrus, SFG=superior frontal gyrus, lOFC=lateral orbitofrontal cortex, ACC=anterior cingulate cortex.

**Figure 2.**
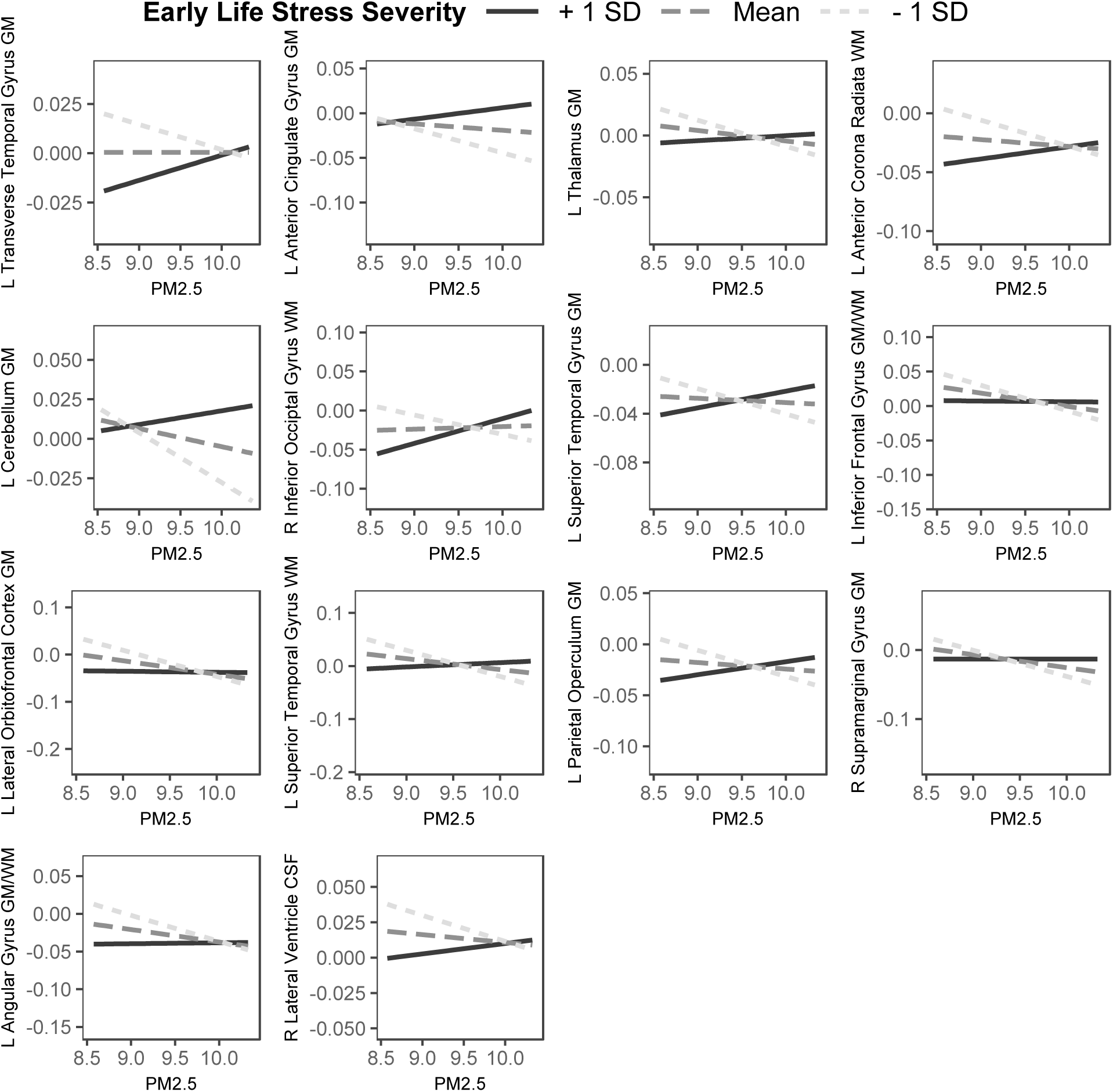
Interactions between early life stress severity and PM2.5 that were positively associated with volumetric regional changes. Note. For the y-axis in each plot, positive and negative values represent expansion and contraction, respectively. L=left; R=right; SLF=superior longitudinal fasciculus; PM2.5= particulate matter < 2.5 micrometers; GM=grey matter; WM=white matter; CSF=cerebrospinal fluid.

**Figure 3.**
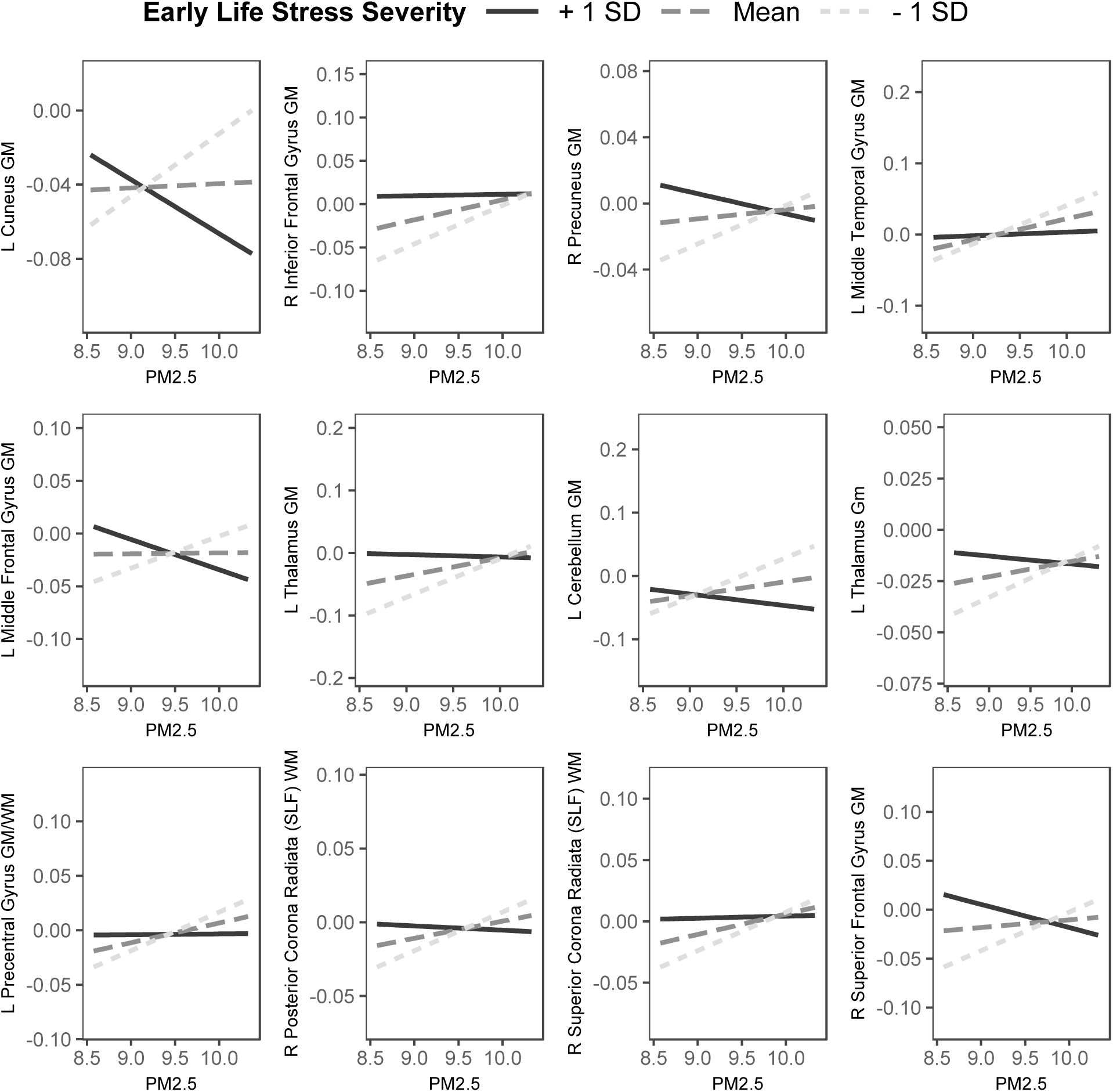
Interactions between early life stress severity and PM2.5 that were negatively associated with volumetric regional changes. Note. For the y-axis in each plot, positive and negative values represent expansion and contraction, respectively. L=left; R=right; SLF=superior longitudinal fasciculus; PM2.5= particulate matter < 2.5 micrometers; GM=grey matter; WM=white matter.

We probed these interactions to determine whether PM2.5 was associated with volumetric changes (i.e., more expansion or contraction) for adolescents who had experienced less, average, or more severe ELS (i.e., less and more severe ELS defined as 1 SD below and above the mean, respectively). Figures 2 and 3 illustrates the estimated positive and negative interaction effects, respectively. End-points for the plotted lines were capped at 1 SD above and below the mean of PM2.5 (34). As Table 2 shows, in 19 of 26 clusters PM2.5 was significantly associated with volumetric changes in adolescents who had experienced less severe ELS (|*β|* ranging from .31 to .59, all *p*<.037); in contrast, PM2.5 was unrelated to volumetric change in adolescents who had experienced more severe ELS (|*β|* ranging from .01 to .18, all *p*>.053). These clusters were located in, among other regions, the inferior frontal gyrus, various temporal and parietal cortical regions, the thalamus, and the corona radiata (in an area overlapping with the superior longitudinal fasciculus). The opposite pattern – that PM2.5 was associated with volumetric change in youth who had experienced more severe ELS and not in youth who had experienced less severe ELS – was obtained in three of the seven remaining clusters (|*β|* ranging from .22 to .28, all *p*<.017). These clusters were located in the left transverse temporal gyrus, left middle frontal gyrus, and the white matter region of the right inferior occipital gyrus. In the remaining four clusters, PM2.5 was associated with volumetric changes at both lower and higher severity of ELS, but in opposite directions. These clusters were located in the left parietal operculum, left superior temporal gyrus, right superior frontal gyrus, and left cuneus.

**Table 2.** Associations between PM2.5 and volumetric change for youth who experienced less and more severe early life stress. Note. Slope estimates represent the association between fine particulate matter (PM2.5) and volumetric regional change (within clusters) at lower (−1 SD) and higher (+1 SD) values of early life stress severity. These estimates are adjusted for age, sex, time interval from time 1 to time 2, total brain volume at time 1, and neighborhood socioeconomic disadvantage. L=left; R=right; SLF=superior longitudinal fasciculus.

| Outcome | Association with<br>PM2.5 at Lower<br>ELS |  | Association with<br>PM2.5 at Higher<br>ELS |  |
| --- | --- | --- | --- | --- |
| | $\beta$ | $p$ | $\beta$ | $p$ |
| **Positive Interactions** |  |  |  |  |
| L Transverse Temporal Gyrus | -.24 | .08 | .22 | .02 |
| L Anterior Cingulate Gyrus | -.31 | .03 | .12 | .18 |
| L Thalamus | -.39 | .008 | .08 | .37 |
| L Anterior Corona Radiata | -.31 | .04 | .15 | .10 |
| L Cerebellum | -.34 | .02 | .08 | .32 |
| R Inferior Occipital Gyrus | -.22 | .12 | .28 | .002 |
| L Superior Temporal Gyrus | -.29 | .04 | .20 | .03 |
| L Inferior Frontal Gyrus | -.59 | <.001 | -.03 | .71 |
| L Lateral Orbitofrontal Cortex | -.49 | <.001 | -.02 | .81 |
| L Superior Temporal Gyrus | -.37 | .02 | .04 | .64 |
| L Parietal Operculum | -.45 | .001 | .20 | .02 |
| R Supramarginal Gyrus | -.51 | <.001 | .01 | .94 |
| L Angular Gyrus | -.53 | <.001 | .01 | .89 |
| R Lateral Ventricle | -.44 | .001 | .16 | .06 |
| **Negative Interactions** |  |  |  |  |
| L Cuneus | .29 | .04 | -.28 | .02 |
| R Inferior Frontal Gyrus | .47 | <.001 | .03 | .76 |
| R Precuneus | .33 | .02 | -.18 | .05 |
| L Middle Temporal Gyrus | .51 | <.001 | .06 | .49 |
| L Middle Frontal Gyrus | .24 | .09 | -.22 | .01 |
| L Thalamus | .46 | .002 | -.04 | .69 |
| L Cerebellum | .38 | .007 | -.12 | .19 |
| L Thalamus | .40 | .005 | -.09 | .32 |
| L Precentral Gyrus | .58 | <.001 | .01 | .91 |
| R Posterior Corona Radiata (SLF) | .46 | .002 | -.06 | .53 |
| R Superior Corona Radiata (SLF) | .45 | .003 | .03 | .78 |
| R Superior Frontal Gyrus | .32 | .02 | -.21 | .02 |

## Discussion

A growing body of work is recognizing the importance of considering how stress and air pollution interact to affect health (19–27). To date, however, this research has mostly focused on respiratory health; in contrast, neuroimaging studies have considered the contributions of psychosocial and environmental risk factors to brain development only independently. Our findings integrate and extend previous studies by showing that the impact of fine particle air pollution (environmental risk factor) on adolescent brain development depends on the severity of ELS, a psychosocial risk factor. Compared to adolescents who had experienced more severe ELS, adolescents with a history of less severe ELS were more sensitive to the effects of PM2.5 in terms of volumetric expansion and contraction across a number of brain regions involved in cognitive and affective functioning, including prefrontal regions, the thalamus, and central white matter. There were far fewer links between PM2.5 and brain development in youth exposed to more severe ELS. These findings suggest that more severe experiences of ELS calibrate or limit neurobiological sensitivity to the effects of PM2.5 on brain development. These findings could also be interpreted as being consistent with a saturation effect model. However, the lack of significant conditional effects of ELS on the observed clusters (after controlling for multiple comparisons) is not consistent with an important prediction of the saturation effect model – that ELS will have a robust effect on brain development that overpowers any additional effect of pollution. Thus, the calibration interpretation is more consistent with our findings than is the saturation effect model.

The observed patterns of regional contraction and expansion related to the interaction of ELS and PM2.5 may reflect cumulative allostatic load related to either psychosocial or environmental stressors. Systemic inflammation related to PM2.5 and/or to particles that directly reach the brain has been posited to promote neuroinflammation and cell death via such processes as oxidative stress, altered brain metabolism, and microglial dysfunction (35–37). Interestingly, ELS affects many of these same biological pathways implicated in apoptosis and neurogenesis, allostatic load, and inflammation (20, 21, 38). In the present study, the brain regions that showed greater contraction, or shrinking, related to severe ELS or increased PM2.5 may be experiencing accelerated cell death. Conversely, brain regions that showed greater expansion related to severe ELS or increased PM2.5 may be undergoing neural reorganization. For example, high levels of ELS or PM2.5 may support the consistent activation and strengthening of pathways that are atypical in the absence of psychosocial and environmental risk factors. Further research is needed to elucidate the specific biological mechanisms that underlie regional expansion and contraction.

Our findings suggest that the experience of more severe ELS minimizes any additional effects of PM2.5 on volumetric expansion and contraction across various grey and white matter brain regions in adolescence. These findings are consistent with the perspective that biological embedding of ELS determines responsivity and openness to the effects of other experiences, including those related to the physical environment (30). Indeed, more severe ELS has been associated with patterns of physiology posited to reflect openness or susceptibility to environmental influence (39, 40). The present study is the first, however, to provide evidence that ELS constrains sensitivity to the effects of air pollution on neurobiology.

Our finding that adolescents exposed to less severe ELS are more susceptible to the effects of PM2.5 on brain development stands in contrast to much of the epidemiological literature on respiratory health, which has largely supported a double exposure model. There are two possible explanations for this inconsistency. First, our study focused on brain development rather than on respiratory health as the outcome of interest. The brain plays a central role in the biological embedding of early social and physical experiences that shape responses and sensitivity to subsequent psychosocial and physical environmental input (29, 30). Thus, the interaction between ELS and PM2.5 may have a different effect on the brain than on other neurobiological systems that are less centrally involved in orchestrating responses to stress and air pollution. Second, the extent to which the interaction effects of ELS and PM2.5 are specific to the overall range of PM2.5 in our study is unclear. It is possible that much higher levels of PM2.5 will affect brain development in youth regardless of ELS history, or that they will change interaction effects such that ELS and PM2.5 will compound risk for effects on brain development.

We found in this study that adolescents who experienced less severe ELS but who lived in neighborhoods with more PM2.5 had patterns of brain development similar to those of adolescents who experienced moderate to severe ELS. This observation is based on many of the interaction plots showing that the endpoint value for the slope at low ELS and high PM2.5 was similar to the endpoint value for the slope at average and high ELS (regardless of PM2.5). For example, in the left lateral orbitofrontal cortex, adolescents exposed to less severe ELS but who lived in more polluted neighborhoods, and adolescents exposed to average to more severe ELS (regardless of air pollution), both demonstrated a reduction in grey matter volume; in contrast, adolescents who had experienced less severe ELS and who lived in less polluted neighborhoods demonstrated grey matter expansion in this region. Thus, even in the absence of experiencing severe ELS, exposure to high levels of PM2.5 appears to have comparable effects on brain development; therefore, more severe ELS and higher PM2.5 may represent different pathways to similar patterns of regional brain contraction and expansion.

Given that many of the brain regions that were associated with PM2.5 are involved in affective and cognitive functioning, our findings have implications for adolescent well-being. For example, the orbitofrontal cortex and thalamus play key roles in emotion-related processes, such as representing the reward value of goals (41, 42). Prefrontal regions such as the inferior and superior frontal gyrus, and central white matter such as the superior longitudinal fasciculus/*corona radiata,* are involved in higher cognitive processes, including executive function and attention (43, 44). TBM does not allow us to separate *corona radiata* from superior longitudinal fasciculus, and future studies using tractography should examine this distinction more explicitly. Furthermore, structural variation in these regions has been implicated in difficulties that emerge during adolescence, including depression, risk-taking, and substance abuse (45–47). Interestingly, recent research has found that increased exposure to PM2.5 is also associated with decreased cognitive development and greater risk for emotional problems (10, 13, 16). Future prospective studies with larger samples should be conducted to test whether volumetric changes in the regions observed in our study might be mechanisms by which PM2.5 affects cognitive, affective, and behavioral functioning in adolescence.

Much of the research to date on air pollution and adolescent brain development has focused on samples in metropolitan areas with higher concentrations of PM2.5 than those documented in our study. Our findings suggest that for adolescents with a history of less severe ELS, the effects of PM2.5 on brain development are evident even at relatively low levels of PM2.5. That said, we should note that the San Francisco Bay Area, from which we recruited our participants, is among the metropolitan areas in the United States with the worst particle air pollution (48). Whether our findings generalize to adolescents living in relatively less air polluted regions of the United States is an open question.

We think that our use of longitudinal TBM is a significant strength of this study. Most studies to date have examined PM2.5-related effects on the brain at a single time point. In contrast, our findings address regional changes in brain structure over time. Certainly, however, it will be important for future research to conduct three or more repeated measures of brain structure or function in order to assess trajectories of brain development that are linked with ELS, PM2.5, and their interaction.

We should note a number of limitations of this study. First, our measure of PM2.5 was based primarily on census tract data obtained from 2012 to 2014. Because we do not know how long adolescents lived at their current addresses, we cannot determine whether our findings are specific to cumulative PM2.5 exposure and/or PM2.5 exposure during a particular developmental window. Nevertheless, we should note that individual differences in air pollution exposure may be stable across development, given that youth tend to grow up in similar neighborhoods over the course of their childhood and adolescence, and that spatial differences in air pollution measurements are stable over long periods of time (49). Thus, differentiating timing from duration effects is a challenge for neighborhood air pollution research in general (50). Second, we measured PM2.5 at the community level. Adolescents living in these communities may vary in their direct exposure to ambient PM2.5 (e.g., spending more time indoor versus outdoor or spending more or less time in the community), as well as in the dose that they receive (e.g., increased physical activity that increases ambient PM2.5 deposition). Although air pollution can vary within a neighborhood based on proximity to highways and local street traffic (51), PM2.5 concentrations do not differ as a function of proximity to highways (52); further, between-neighborhood variability in PM2.5 has been shown to be greater than within-neighborhood variability (53). Taken together, these studies support our focus on between-neighborhood variation in PM2.5. Nevertheless, capturing individual differences in exposure and dose of PM2.5 will be important for determining the degree to which effects are specific to this form of air pollution. As a related point, our observational findings do not identify the mechanisms by which increased PM2.5 may lead to regional brain expansion and contraction, and do not causally link PM2.5 to brain development. Although it is likely that PM2.5 affects the developing brain directly, it is also possible that our observed effects are due to unmeasured factors related to air pollution (e.g., traffic-related noise pollution). We controlled for variables that have been posited to be correlated with ambient air pollution exposure, including minority status, poverty rate, educational attainment, and unemployment (19–22) (see supplementary materials), but we cannot definitively rule out all possible confounds or selection variables that might lead individuals to reside in certain neighborhoods that have more or less air pollution (54). In this context, however, PM2.5 was not associated with other variables in our model, which suggests that neighborhood-level PM2.5 was not a proxy for race/ethnicity or socioeconomic status in our sample. Nevertheless, randomized trials are necessary to determine whether changing PM2.5 exposure affects brain measures differently for youth exposed to more versus less severe ELS. Finally, our measure of severity of ELS was based on retrospective self-report, which may be subject to biased reporting, although in this study we used raters’ coding of objective ELS severity (see below).

Despite these limitations, this study is the first to provide evidence that psychosocial risk factors moderate the impact of environmental risk factors on adolescent brain development. These effects were remarkably consistent across different brain regions and suggest that PM2.5 is associated with volumetric expansion and contraction for adolescents who are exposed to less severe ELS. These findings provide a nuanced, rich understanding of which adolescents may be most sensitive to PM2.5 in terms of brain development, and serve as a call for more research and theory that considers the joint effects of air pollution and ELS on the brain. In addition, for youth exposed to less severe ELS, despite living in neighborhoods with PM2.5 levels that are considered safe by the EPA, elevated PM2.5 was nevertheless associated with patterns of regional volumetric expansion and contraction that were comparable to those observed in adolescents with more severe ELS experiences. Although ELS clearly disrupts the developing brain, increased PM2.5 may produce similar adverse effects even in the absence of a history of severe ELS.

## Methods

### Sample

Participants were healthy English-speaking adolescents living in the San Francisco and San Jose Bay Area. Boys and girls were matched on pubertal stage at study entry. Given that girls typically reach sexual maturity earlier than boys, girls were slightly younger than boys in our sample (mean difference=0.69 years, *t*(114)=3.61, p<.001). Most of the data for the current analyses were collected from 2013-2018. Our PM2.5 measure was based on census tract data collected from 2012-2014 (see below). We used participants’ mailing addresses to identify census tracts. Adolescents and their parents signed assent and consent forms, respectively. The Stanford University Institutional Review Board approved the study protocol.

### PM2.5

PM2.5 concentrations at the census tract level were assessed using air quality monitoring station data collected by the California Air Resources Board and compiled and made publicly available by the California Office of Environmental Health Hazard Assessment (OEHHA; 55). Ordinary kriging was used to estimate average PM2.5 concentration values at the geographic center of each census tract from nearby monitors. Quarterly estimates were averaged to compute annual means, which were then averaged over the three year period from 2012-2014.

### Early Life Stress

At Time 1, adolescents completed an interview to asses exposure to 30 different types of stressful experiences using a modified version of the Traumatic Events Screening Inventory for Children (e.g., parental divorce, family mental illness/substance abuse, accidents, etc.) (56). A panel of coders, who were blind to the adolescents’ reactions during the interview, rated interview responses for objective severity using a modified version of the UCLA Life Stress Interview coding system (57); the objective severity of each reported stressful event was rated on scale ranging from 1 (non-event or no impact) to 5 (extremely severe impact) with half-point increments. Ratings were summed across stressful events that were rated at 1.5 or higher to create a cumulative measure of ELS severity for each adolescent.

### Neighborhood Socioeconomic Disadvantage

Neighborhood-level disadvantage was assessed using poverty, educational attainment, unemployment, and housing burden data from multiple nationally representative datasets (e.g., The American Community Survey). The OEHHA converted these data into percentiles representing the amount of socioeconomic disadvantage in a given census tract relative to other census tracts in California. The percentiles for poverty, educational attainment, unemployment, and housing burden were averaged to create a measure of overall neighborhood socioeconomic disadvantage.

### Scan

MRI scans were acquired at the Center for Cognitive and Neurobiological Imaging at Stanford University using a 3 T Discovery MR750 (GE Medical Systems, Milwaukee, WI, USA) equipped with a 32-channel head coil (Nova Medical, Wilmington, MA, USA). Whole-brain T1-weighted images (T1w) were collected using the following spoiled gradient echo pulse sequence: 186 sagittal slices; TR (repetition time)/TE (echo time)/TI (inversion time) = 6.24/2.34/450ms; flip angle = 12°; voxel size = 0.9 mm×0.9 mm×0.9 mm; scan duration = 5:15 minutes.

### Tensor-Based Morphometry

At both time points, each participant’s T1-weighted anatomical data were N3-corrected using c3d (http://www.itksnap.org) to correct for intensity inhomogeneities. Volumes were automatically skull-stripped using Brainsuite and brain masks were manually edited to remove extraneous skull or meninges by trained neuroanatomical experts (LS, AC, and AO, see Acknowledgements). We used *flirt* (http://fsl.fmrib.ox.ac.uk) to linearly register each participant to a study-specific template. We used a study-specific registration template to obtain the strongest possible registration results (58). We chose a female aged 11 years 5 months (the average for the sample subset included) with a visually normal T1-weighted scan to initialize the linear registration. This exemplar participant was registered to the ICBM template using *flirt*, using 7 degrees of freedom registration with trilinear interpolation, and using mutual information as the similarity function for alignment. Following this, each participant’s masked, N3-corrected T1-weighted image was registered to the participant-template using iterative 6-, 7-, and 9-DOF (degrees of freedom) registration. We concatenated transformation files so that only one resampling step was run. This protocol was modified for this dataset from the original protocol (59, 60). Thirty participants, selected to be representative of the population, were used to make the minimal deformation target (MDT). This representative group was chosen semi-randomly by dividing the sample into tertiles based on exposure to early life stress and randomly selecting 5 females and 5 males from each tertile. The MDT is the template that deviates least from the anatomy of the participants with respect to a mathematically defined metric of difference; in some circumstances, using a MDT can improve statistical power (61). The MDT serves as an unbiased registration target for nonlinear registrations.

Each participant’s masked, non-uniformity-corrected, template-aligned T1-weighted image from the follow-up scan was non-linearly aligned to their first scan, using Advanced Normalization Tools’ Symmetric Normalization (SyN) (62). SyN registration used a multi-level approach, i.e., the “moving” and fixed T1-weighted images were successively less smoothed at each level, with a full resolution registration occurring at the final level. We used 150, 80, 50, and 10 iterations at each level, with a Gaussian kernel smoothing sigma set to 3, 2, 1, and 0 respectively (7.05, 4.7, 2.35, and 0 voxels full width at half maximum), and shrink factors of 4, 2, 2, and 1 respectively. Image similarity was measured using the ANTs implementation of mutual information (63). Image intensities were winsorized, excluding top and bottom 1% of voxels, and histogram matching was used. The output jacobian determinant image showed the direction and magnitude of the change between the participant’s Time 1 and Time 2 anatomical images. Output was visually checked for quality.

#### Statistical Analyses

In our voxel-wise linear regression testing for group differences, we included intracranial volume (ICV) as a covariate. The 9-DOF linear registration that is part of our processing protocol accounts for differences in overall brain scale, removing much of the effect of ICV; nevertheless, we included ICV, computed from the linearly registered image, as a covariate. In a prior analysis of this dataset, we found minimal differences between models with and without ICV as a covariate (64). To test associations with ELS and pollution exposure, we examined several models:

Model (interaction between ELS and PM2.5): *X* ∼ *A* + *β*_*Sex*_*Sex* + *β*_*ageT*1_*AgeT*1 + *β*_*Interval*_*Interval* +*β*_*SES*_*SES* +*β*_*ICVT*1_*ICVT*1 +*β*_*ELS*_*ELS* + *β*_*PM*_*PM*2.5 +*β*_*ELS x PM*_*ELS x PM*2.5 + *ε*
Model (associations with ELS) *X* ∼ *A* + *β*_*Sex*_*Sex* + *β*_*ageT*1_*AgeT*1 + *β*_*Interval*_*Interval* +*β*_*SES*_*SES* + *β*_*ICVT*1_*ICVT*1 +*β*_*ELS*_*ELS* + *ε*
Model (associations with PM2.5) *X* ∼ *A* + *β*_*Sex*_*Sex* + *β*_*ageT*1_*AgeT*1 + *β*_*Interval*_*Interval* +*β*_*SES*_*SES* + *β*_*ICVT*1_*ICVT*1 +*β*_*PM*_*PM*2.5 + *ε*

where *X* is the Jacobian determinant value at a given position, *A* is the constant Jacobian determinant term, the βs are the covariate regression coefficients, and ε is an error term. All predictors were centered prior to forming the interaction term. For each model, results were corrected for multiple comparisons across all voxels tested using Searchlight FDR (*q*<0.05) (65). We only ran further analyses and report on those clusters that exceeded our criteria of 50 voxels. Covariates across the models included age, sex, ICV, interval between T1 and T2, and neighborhood-level socioeconomic disadvantage (see supplementary material for sensitivity analyses).

## Acknowledgments

This research was supported by the National Institute of Mental Health Grants R37-MH101495 to IHG, and MH019908 to Allan L. Reiss (funding JGM). We thank the research staff who made this work possible, and the families who participated in this study.

