## supplemental material for "Severity of Early Life Stress Moderates the Effect of Fine Particle Air Pollution on Adolescent Brain Development"

Supplementary Material

Descriptive statistics and zero-order correlations for the main predictor variables and covariates are presented in Table S1.

Prior to probing the ELS by PM2.5 interaction effects, values representing change in brain volume were extracted from each cluster and were inspected for extreme outlier values using boxplots (values greater or less than the mean by 3 times the interquartile range). In clusters containing extreme outliers, regression analyses were rerun after removing these values (no more than 2 values removed in any given cluster), which resulted in 11 clusters no longer being associated with the ELS by PM2.5 interaction. Given that the previously observed interaction effects for these clusters were driven by extreme outlier TBM values, we did not include these clusters in further analyses. An additional cluster was dropped due to the effects of PM2.5 on volumetric change being not statistically different from zero at high and low ELS severity (both $p>.054$). Thus, the final analyses included 26 clusters (see Table 2 in main text).

We performed sensitivity analyses to assess whether the results were impacted by which covariates were included or excluded from the model. These analyses consisted of testing six models that differed from our original model (see main text) in terms of including minority status as a covariate, dropping T1 total brain volume from the model, and separating our socioeconomic disadvantage composite variable into separate components for poverty, education, and unemployment (see Tables S2-S7). More specifically, Table S2 shows the results for a model that includes minority status as an additional covariate. This model also separates socioeconomic disadvantage into separate components. Table S3 shows the results after excluding total brain volume at time 1 from the model. Table S4 shows the results after excluding total brain volume and minority status from the model. Table S5 shows the results after excluding total brain volume, poverty, and unemployment. Table S6 shows the results after excluding total brain volume, education, and unemployment. Table S7 shows the results after excluding total brain volume, education, and poverty. The clusters that were associated with the

ELS by PM2.5 interaction were largely observed in the same brain regions across models; there were some differences in cluster size. Thus, there are minimal differences in the clusters associated with the ELS by PM2.5 interaction depending on which covariates are included in the model.

Table S1. Descriptive Statistics and Correlations for Early Life Stress Severity, Fine Particulate Matter (PM2.5), and Covariates.

| | 1 | 2 | 3 | 4 | 5 | 6 | 7 |
|---|---|---|---|---|---|---|---|
| 1. Sex (male=1) | 1 | | | | | | |
| 2. T1 Age | .32*** | 1 | | | | | |
| 3. T1-T2 Interval | -.11 | -.03 | 1 | | | | |
| 4. T1 Total Brain Volume | -.06 | .05 | -.09 | 1 | | | |
| 5. Neighborhood Disadvantage (% ranking) | -.00 | -.18* | -.08 | .01 | 1 | | |
| 6. ELS Severity | -.00 | .01 | .04 | .11 | .10 | 1 | |
| 7. PM2.5 (µg/m$^3$) | -.14 | -.11 | -.12 | .11 | -.07 | -.08 | 1 |
| Mean or % | 44% | 11.50 | 1.96 | 1347.34 | 28.41 | 6.65 | 9.46 |
| SD | | 1.08 | 0.45 | 14.97 | 18.48 | 5.20 | 0.91 |
| Range | | 9.35-13.95 | 1.18-4.16 | 1298.14-1405.08 | .75-79.50 | .00-24.00 | 7.02-13.44 |

*Note*. ***p<.001, *p<.05. T1=Time 1; T2=Time2; ELS=Early Life Stress; PM2.5=particulate matter < 2.5 micrometers. Females were coded as 0 and males were coded as 1.

Table S2. Clusters showing a significant interaction between ELS and PM on regional brain volume. This model contains the following covariates: T1 total brain volume, sex, age, interval between T1 and T2, ELS, PM2.5, minority status, and education, poverty, and unemployment as separate predictors (instead of the socioeconomic disadvantage composite).

**Positive Interactions**

| Voxels | Max. T-stat | X | Y | Z | Side | Structure | Tissue |
|---|---|---|---|---|---|---|---|
| 752 | 4.8 | 22 | -34 | 18 | R | Lateral ventricle | CSF |
| 350 | 4.03 | -44 | -44 | 23 | L | Angular gyrus | GM/WM |
| 168 | 4.17 | -22 | 12 | -21 | L | Lateral orbitofrontal cortex | GM |
| 163 | 3.92 | 39 | -40 | 45 | R | Supramarginal gyrus | GM |
| 150 | 5.39 | -47 | 11 | -14 | L | Superior temporal gyrus | WM |
| 117 | 4.13 | -48 | -33 | 17 | L | Parietal operculum | GM |
| 114 | 4.76 | -44 | 5 | 9 | L | Inferior frontal gyrus | GM/WM |
| 105 | 4.65 | -27 | -76 | -46 | L | Cerebellum | GM |
| 90 | 4.33 | 54 | -65 | 9 | R | Inferior occipital gyrus | WM |
| 88 | 4.95 | -59 | -58 | -10 | L | Inferior temporal gyrus | GM |
| 86 | 4.41 | -45 | -41 | 5 | L | Superior temporal gyrus | GM |
| 85 | 5.17 | 63 | -3 | 30 | R | Postcentral gyrus | GM |
| 83 | 4.17 | -45 | -71 | 5 | L | Lateral occipital gyrus | GM |
| 81 | 4.35 | 32 | -24 | 58 | R | Precentral gyrus | GM |
| 63 | 3.57 | 25 | -46 | 13 | R | Splenium | WM |

**Negative Interactions**

| Voxels | Max. T-stat | X | Y | Z | Side | Structure | Tissue |
|---|---|---|---|---|---|---|---|
| 1205 | 4.46 | 6 | 37 | 51 | R | Superior frontal gyrus | GM |
| 448 | 3.67 | 30 | -22 | 23 | R | Superior corona radiata (SLF) | WM |
| 273 | 4 | -8 | -22 | 8 | L | Thalamus | GM |
| 270 | 4.03 | -31 | -54 | -57 | L | Cerebellum | GM |
| 248 | 4.17 | -39 | -15 | 37 | L | Precentral gyrus | GM/WM |
| 218 | 4.81 | -10 | -22 | 70 | L | Thalamus | GM |
| 188 | 3.79 | 35 | -39 | 16 | R | Posterior corona radiata (SLF) | WM |
| 166 | 4.78 | -32 | 4 | 59 | L | Middle frontal gyrus | GM |
| 152 | 5.52 | -14 | -98 | -10 | L | Cuneus | GM |
| 123 | 5.35 | 10 | -62 | 59 | R | Precuneus | GM |
| 122 | 7.56 | -20 | -64 | 56 | L | Superior parietal lobule | GM |
| 117 | 5.61 | 34 | 26 | -19 | R | Lateral orbitofrontal cortex | GM |
| 112 | 4.24 | -41 | 50 | 18 | L | Middle frontal gyrus | GM |
| 81 | 4.06 | -41 | 54 | 1 | L | Middle frontal gyrus | GM |
| 81 | 4.48 | -5 | 36 | -10 | L | Medial orbitofrontal cortex | GM |
| 72 | 4.36 | 25 | 64 | 12 | R | Superior frontal gyrus | GM |
| 69 | 3.92 | -42 | 14 | -38 | L | Middle temporal gyrus | GM |
| 68 | 3.91 | -8 | -64 | 4 | L | Lingual gyrus | GM |
| 64 | 3.63 | 17 | -79 | -47 | R | Cerebellum | GM |
| 58 | 3.69 | 51 | 12 | 23 | R | Inferior frontal gyrus | GM |
| 57 | 4.01 | 10 | -58 | 40 | R | Precuneus | GM |
| 54 | 4.43 | -25 | -97 | -4 | L | Cuneus | GM |
| 52 | 3.97 | 30 | -43 | 48 | R | Superior parietal lobule | WM |

Table S3. Clusters showing a significant interaction between ELS and PM on regional brain volume. This model contains the covariates from the Table S2 model excluding T1 total brain volume.

**Positive Interactions**

| Voxels | Max. T-stat | X | Y | Z | Side | Structure | Tissue |
|---|---|---|---|---|---|---|---|
| 703 | 4.83 | 22 | -34 | 18 | R | Lateral ventricle | CSF |
| 321 | 4.06 | -44 | -44 | 23 | L | Angular gyrus | GM/WM |
| 172 | 3.95 | 39 | -40 | 45 | R | Supramarginal gyrus | GM |
| 166 | 4.2 | -22 | 12 | -21 | L | Lateral orbitofrontal cortex | GM |
| 144 | 5.37 | -47 | 11 | -14 | L | Superior temporal gyrus | WM |
| 114 | 4.13 | -48 | -33 | 17 | L | Parietal operculum | GM |
| 110 | 4.75 | -44 | 5 | 9 | L | Inferior frontal gyrus | GM/WM |
| 107 | 4.62 | -27 | -76 | -46 | L | Cerebellum | GM |
| 92 | 4.35 | 54 | -65 | 9 | R | Inferior occipital gyrus | WM |
| 89 | 4.93 | -59 | -58 | -10 | L | Inferior temporal gyrus | GM |
| 88 | 4.44 | -45 | -41 | 5 | L | Superior temporal gyrus | GM |
| 85 | 5.06 | 63 | -3 | 30 | R | Postcentral gyrus | GM |
| 80 | 4.36 | 32 | -24 | 58 | R | Precentral gyrus | GM |
| 79 | 4 | -46 | -71 | 5 | L | Lateral occipital gyrus | GM |

**Negative Interactions**

| Voxels | Max. T-stat | X | Y | Z | Side | Structure | Tissue |
|---|---|---|---|---|---|---|---|
| 1262 | 4.45 | -17 | 21 | 57 | L | Superior frontal gyrus | GM |
| 461 | 3.69 | 30 | -22 | 23 | R | Superior corona radiata (SLF) | WM |
| 280 | 4.03 | -8 | -22 | 8 | L | Thalamus | GM |
| 272 | 4.01 | -31 | -54 | -57 | L | Cerebellum | GM |
| 248 | 4.18 | -39 | -15 | 37 | L | Precentral gyrus | GM/WM |
| 221 | 4.84 | -10 | -22 | 70 | L | Thalamus | GM |
| 207 | 3.82 | 35 | -39 | 16 | R | Posterior corona radiata (SLF) | WM |
| 171 | 4.81 | -32 | 4 | 59 | L | Middle frontal gyrus | GM |
| 153 | 5.54 | -14 | -98 | -10 | L | Cuneus | GM |
| 126 | 5.38 | 10 | -62 | 59 | R | Precuneus | GM |
| 122 | 7.58 | -20 | -64 | 56 | L | Superior parietal lobule | GM |
| 121 | 5.59 | 34 | 26 | -19 | R | Lateral orbitofrontal cortex | GM |
| 118 | 4.26 | -41 | 50 | 18 | L | Middle frontal gyrus | GM |
| 84 | 4.07 | -41 | 54 | 1 | L | Middle frontal gyrus | GM |
| 78 | 4.49 | -5 | 36 | -10 | L | Medial orbitofrontal cortex | GM |
| 73 | 4.39 | 25 | 64 | 12 | R | Superior frontal gyrus | GM |
| 67 | 3.89 | -8 | -64 | 4 | L | Lingual gyrus | GM |
| 66 | 3.64 | 17 | -79 | -47 | R | Cerebellum | GM |
| 59 | 4.03 | 10 | -58 | 40 | R | Precuneus | GM |
| 55 | 4.46 | -25 | -97 | -4 | L | Cuneus | GM |
| 53 | 3.78 | -42 | 14 | -38 | L | Middle temporal gyrus | GM |
| 52 | 3.97 | 30 | -43 | 48 | R | Superior parietal lobule | WM |

Table S4. Clusters showing a significant interaction between ELS and PM on regional brain volume. This model contains the covariates from the Table S3 model except for minority status.

**Positive Interactions**

| Voxels | Max. T-stat | X | Y | Z | Side | Structure | Tissue |
|---|---|---|---|---|---|---|---|
| 727 | 4.86 | 22 | -34 | 18 | R | Lateral ventricle | CSF |
| 333 | 4.08 | -44 | -44 | 23 | L | Angular gyrus | GM/WM |
| 181 | 3.97 | 39 | -40 | 45 | R | Supramarginal gyrus | GM |
| 174 | 4.23 | -22 | 12 | -21 | L | Lateral orbitofrontal cortex | GM |
| 146 | 5.4 | -47 | 11 | -14 | L | Superior temporal gyrus | WM |
| 110 | 4.74 | -44 | 5 | 9 | L | Inferior frontal gyrus | GM/WM |
| 109 | 4.1 | -48 | -33 | 17 | L | Parietal operculum | GM |
| 109 | 4.63 | -26 | -76 | -46 | L | Cerebellum | GM |
| 96 | 4.36 | 54 | -65 | 9 | R | Inferior occipital gyrus | WM |
| 90 | 4.46 | -45 | -41 | 5 | L | Superior temporal gyrus | GM |
| 89 | 4.95 | -59 | -58 | -10 | L | Inferior temporal gyrus | GM |
| 84 | 5.08 | 63 | -3 | 30 | R | Postcentral gyrus | GM |
| 81 | 4.38 | 32 | -24 | 58 | R | Precentral gyrus | GM |
| 80 | 4.02 | -46 | -71 | 5 | L | Lateral occipital gyrus | GM |

**Negative Interactions**

| Voxels | Max. T-stat | X | Y | Z | Side | Structure | Tissue |
|---|---|---|---|---|---|---|---|
| 1274 | 4.43 | -16 | 21 | 57 | L | Superior frontal gyrus | GM |
| 479 | 3.71 | 30 | -22 | 23 | R | Superior corona radiata (SLF) | WM |
| 285 | 4.05 | -8 | -22 | 8 | L | Thalamus | GM |
| 280 | 4.03 | -31 | -54 | -57 | L | Cerebellum | GM |
| 253 | 4.2 | -39 | -15 | 37 | L | Precentral gyrus | GM/WM |
| 223 | 4.64 | -10 | -22 | 70 | L | Thalamus | GM |
| 220 | 3.84 | 35 | -39 | 16 | R | Posterior corona radiata (SLF) | WM |
| 165 | 4.83 | -32 | 4 | 59 | L | Middle frontal gyrus | GM |
| 146 | 5.55 | -14 | -98 | -10 | L | Cuneus | GM |
| 126 | 5.41 | 10 | -62 | 59 | R | Precuneus | GM |
| 125 | 7.62 | -20 | -64 | 56 | L | Superior parietal lobule | GM |
| 125 | 4.27 | -41 | 50 | 18 | L | Middle frontal gyrus | GM |
| 122 | 5.51 | 34 | 26 | -19 | R | Lateral orbitofrontal cortex | GM |
| 85 | 4.09 | -41 | 54 | 1 | L | Middle frontal gyrus | GM |
| 81 | 4.51 | -5 | 36 | -10 | L | Medial orbitofrontal cortex | GM |
| 69 | 3.92 | -8 | -64 | 4 | L | Lingual gyrus | GM |
| 67 | 3.63 | 16 | -75 | -48 | R | Cerebellum | GM |
| 63 | 4.21 | 25 | 64 | 12 | R | Superior frontal gyrus | GM |
| 60 | 4.04 | 10 | -58 | 40 | R | Precuneus | GM |
| 60 | 3.8 | -42 | 14 | -38 | L | Middle temporal gyrus | GM |
| 55 | 4.45 | -25 | -97 | -4 | L | Cuneus | GM |
| 53 | 3.99 | 30 | -43 | 48 | R | Superior parietal lobule | WM |

Table S5. Clusters showing a significant interaction between ELS and PM on regional brain volume. This model contains the covariates from the Table S3 model but only includes education, not poverty or unemployment.

**Positive Interactions**

| Voxels | Max. T-stat | X | Y | Z | Side | Structure | Tissue |
|---|---|---|---|---|---|---|---|
| 781 | 4.95 | 22 | -34 | 18 | R | Lateral ventricle | CSF |
| 337 | 4.07 | -44 | -44 | 23 | L | Angular gyrus | GM/WM |
| 149 | 5.62 | -47 | 11 | -14 | L | Superior temporal gyrus | WM |
| 129 | 3.86 | 39 | -40 | 45 | R | Supramarginal gyrus | GM |
| 128 | 4.17 | -48 | -33 | 17 | L | Parietal operculum | GM |
| 119 | 3.87 | -22 | 12 | -21 | L | Lateral orbitofrontal cortex | GM |
| 108 | 4.61 | -44 | 5 | 9 | L | Inferior frontal gyrus | GM/WM |
| 98 | 4.65 | -45 | -41 | 5 | L | Superior temporal gyrus | GM |
| 96 | 5.29 | 63 | -3 | 30 | R | Postcentral gyrus | GM |
| 89 | 4.27 | 54 | -65 | 9 | R | Inferior occipital gyrus | WM |
| 85 | 5 | -59 | -59 | -10 | L | Inferior temporal gyrus | GM |
| 85 | 4.68 | -27 | -76 | -46 | L | Cerebellum | GM |
| 79 | 4.38 | 32 | -24 | 58 | R | Precentral gyrus | GM |
| 56 | 3.83 | -17 | -25 | 14 | L | Thalamus | GM |
| 51 | 3.58 | -9 | 29 | 2 | L | Genu | WM |

**Negative Interactions**

| Voxels | Max. T-stat | X | Y | Z | Side | Structure | Tissue |
|---|---|---|---|---|---|---|---|
| 1269 | 4.39 | -6 | 28 | 57 | L | Superior frontal gyrus | GM |
| 406 | 3.64 | 30 | -22 | 23 | R | Superior corona radiata (SLF) | WM |
| 297 | 3.97 | 35 | -39 | 16 | R | Posterior corona radiata (SLF) | WM |
| 284 | 4.17 | -7 | -24 | 8 | L | Thalamus | GM |
| 258 | 4.28 | -39 | -15 | 37 | L | Precentral gyrus | GM/WM |
| 235 | 4.94 | -10 | -22 | 70 | L | Thalamus | GM |
| 172 | 5.64 | -14 | -98 | -10 | L | Cuneus | GM |
| 140 | 7.76 | -20 | -64 | 56 | L | Superior parietal lobule | GM |
| 137 | 5.51 | 10 | -62 | 59 | R | Precuneus | GM |
| 135 | 5.83 | 34 | 26 | -19 | R | Lateral orbitofrontal cortex | GM |
| 129 | 4.31 | -40 | 42 | 20 | L | Middle frontal gyrus | GM |
| 99 | 4.38 | -32 | 4 | 59 | L | Middle frontal gyrus | GM |
| 94 | 4.66 | -5 | 36 | -10 | L | Medial orbitofrontal cortex | GM |
| 83 | 3.61 | 52 | 13 | 24 | R | Inferior frontal gyrus | GM |
| 83 | 3.86 | 17 | -79 | -47 | R | Cerebellum | GM |
| 74 | 4.11 | 10 | -58 | 40 | R | Precuneus | GM |
| 73 | 4.05 | -41 | 54 | 1 | L | Middle frontal gyrus | GM |
| 71 | 4.05 | 30 | -42 | 49 | R | Superior parietal lobule | WM |
| 69 | 4.44 | 25 | 64 | 12 | R | Superior frontal gyrus | GM |
| 57 | 3.82 | -43 | 14 | -38 | L | Middle temporal gyrus | GM |
| 52 | 4.43 | -26 | -97 | -4 | L | Cuneus | GM |

Table S6. Clusters showing a significant interaction between ELS and PM on regional brain volume. This model contains the covariates from the Table S3 model but only includes poverty, not education or unemployment.

**Positive Interactions**

| Voxels | Max. T-stat | X | Y | Z | Side | Structure | Tissue |
|---|---|---|---|---|---|---|---|
| 889 | 4.8 | 22 | -34 | 18 | R | Lateral ventricle | CSF |
| 361 | 4.09 | -44 | -44 | 23 | L | Angular gyrus | GM/WM |
| 234 | 3.97 | 39 | -40 | 45 | R | Supramarginal gyrus | GM |
| 199 | 4.32 | -22 | 12 | -21 | L | Lateral orbitofrontal cortex | GM |
| 149 | 5.45 | -47 | 10 | -14 | L | Superior temporal gyrus | WM |
| 133 | 4.86 | -44 | 5 | 9 | L | Inferior frontal gyrus | GM/WM |
| 127 | 4.2 | -48 | -33 | 17 | L | Parietal operculum | GM |
| 118 | 4.47 | 54 | -65 | 9 | R | Inferior occipital gyrus | WM |
| 116 | 4.66 | -27 | -76 | -46 | L | Cerebellum | GM |
| 89 | 4.42 | -45 | -42 | 5 | L | Superior temporal gyrus | GM |
| 87 | 4.98 | -59 | -58 | -10 | L | Inferior temporal gyrus | GM |
| 85 | 5.02 | 63 | -3 | 31 | R | Postcentral gyrus | GM |
| 82 | 3.96 | -46 | -71 | 5 | L | Lateral occipital gyrus | GM |
| 62 | 4.12 | 32 | -24 | 58 | R | Precentral gyrus | GM |
| **Negative Interactions** | | | | | | | |
| **Voxels** | **Max. T-stat** | **X** | **Y** | **Z** | **Side** | **Structure** | **Tissue** |
| 1294 | 4.45 | 6 | 37 | 51 | R | Superior frontal gyrus | GM |
| 578 | 3.8 | 30 | -22 | 23 | R | Superior corona radiata (SLF) | WM |
| 323 | 4.13 | -31 | -54 | -57 | L | Cerebellum | GM |
| 295 | 4.11 | -8 | -22 | 8 | L | Thalamus | GM |
| 256 | 4.17 | -39 | -15 | 37 | L | Precentral gyrus | GM/WM |
| 253 | 3.87 | 35 | -39 | 16 | R | Posterior corona radiata (SLF) | WM |
| 189 | 4.64 | -10 | -22 | 70 | L | Thalamus | GM |
| 166 | 5.5 | -14 | -98 | -10 | L | Cuneus | GM |
| 159 | 3.67 | -5 | -68 | -42 | L | Cerebellum | GM |
| 158 | 4.84 | -32 | 4 | 59 | L | Middle frontal gyrus | GM |
| 133 | 4.36 | -41 | 50 | 18 | L | Middle frontal gyrus | GM |
| 126 | 5.58 | 34 | 26 | -19 | R | Lateral orbitofrontal cortex | GM |
| 125 | 5.24 | 11 | -62 | 60 | R | Precuneus | GM |
| 108 | 7.68 | -20 | -64 | 56 | L | Superior parietal lobule | GM |
| 80 | 4.43 | 25 | 64 | 12 | R | Superior frontal gyrus | GM |
| 78 | 4.46 | -5 | 36 | -10 | L | Medial orbitofrontal cortex | GM |
| 73 | 3.96 | -8 | -64 | 4 | L | Lingual gyrus | GM |
| 62 | 3.84 | -41 | 54 | 1 | L | Middle frontal gyrus | GM |
| 62 | 3.84 | -42 | 14 | -38 | L | Middle temporal gyrus | GM |
| 61 | 3.65 | 16 | -75 | -48 | R | Cerebellum | GM |
| 58 | 3.97 | 10 | -58 | 40 | R | Precuneus | GM |
| 57 | 4.04 | 30 | -43 | 48 | R | Superior parietal lobule | WM |
| 55 | 4.53 | -25 | -97 | -4 | L | Cuneus | GM |
| 51 | 3.74 | -49 | -11 | 6 | L | Planum polare | GM |

Table S7. Clusters showing a significant interaction between ELS and PM on regional brain volume. This model contains the covariates from the Table S3 model but only includes unemployment, not education or poverty.

| | | | | | | | |
|---|---|---|---|---|---|---|---|
| | | | | **Positive Interactions** | | | |
| **Voxels** | **Max. T-stat** | **X** | **Y** | **Z** | **Side** | **Structure** | **Tissue** |
| 1185 | 5 | 22 | -34 | 18 | R | Lateral ventricle | CSF |
| 347 | 4.25 | -45 | -44 | 22 | L | Angular gyrus | GM/WM |
| 236 | 3.67 | 40 | -35 | 42 | R | Supramarginal gyrus | GM |
| 181 | 4.19 | -22 | 12 | -21 | L | Lateral orbitofrontal cortex | GM |
| 179 | 6.03 | -47 | 11 | -14 | L | Superior temporal gyrus | WM |
| 168 | 4.33 | -48 | -33 | 18 | L | Parietal operculum | GM |
| 161 | 5.18 | -27 | -76 | -46 | L | Cerebellum | GM |
| 155 | 4.62 | 54 | -65 | 9 | R | Inferior occipital gyrus | WM |
| 155 | 5.05 | -44 | 5 | 9 | L | Inferior frontal gyrus | GM/WM |
| 140 | 4.11 | -40 | -29 | 6 | L | Transverse temporal gyrus | GM |
| 128 | 4.86 | -45 | -41 | 5 | L | Superior temporal gyrus | GM |
| 126 | 4.11 | -18 | -25 | 13 | L | Thalamus | GM |
| 105 | 5.49 | 63 | -3 | 31 | R | Postcentral gyrus | GM |
| 88 | 4.11 | -23 | 27 | -5 | L | Anterior corona radiata | WM |
| 83 | 4.17 | -10 | 44 | -4 | L | Anterior cingulate gyrus | GM |
| 81 | 5.08 | -59 | -59 | -10 | L | Inferior temporal gyrus | GM |
| 76 | 3.6 | -35 | 25 | -28 | L | Temporal pole | GM |
| 58 | 3.84 | -30 | -46 | -40 | L | Cerebellum | WM |
| 51 | 3.75 | -10 | 29 | 0 | L | Genu | WM |
| | | | | **Negative Interactions** | | | |
| **Voxels** | **Max. T-stat** | **X** | **Y** | **Z** | **Side** | **Structure** | **Tissue** |
| 1159 | 4.58 | 6 | 37 | 51 | R | Superior frontal gyrus | GM |
| 612 | 3.93 | 30 | -22 | 23 | R | Superior corona radiata (SLF) | WM |
| 445 | 4.2 | 35 | -39 | 16 | R | Posterior corona radiata (SLF) | WM |
| 381 | 4.41 | -7 | -24 | 8 | L | Thalamus | GM |
| 361 | 4.17 | -31 | -54 | -57 | L | Cerebellum | GM |
| 283 | 4.15 | -39 | -15 | 37 | L | Precentral gyrus | GM/WM |
| 208 | 5.87 | -14 | -98 | -10 | L | Cuneus | GM |
| 192 | 4.62 | -41 | 50 | 18 | L | Middle frontal gyrus | GM |
| 174 | 5.98 | 34 | 26 | -19 | R | Lateral orbitofrontal cortex | GM |
| 167 | 4.54 | -10 | -22 | 70 | L | Thalamus | GM |
| 151 | 3.45 | 6 | -20 | 5 | R | Thalamus | GM |
| 149 | 5.27 | 11 | -62 | 60 | R | Precuneus | GM |
| 127 | 8.03 | -20 | -64 | 56 | L | Superior parietal lobule | GM |
| 120 | 4.45 | -32 | 4 | 59 | L | Middle frontal gyrus | GM |
| 104 | 3.97 | 16 | -75 | -48 | R | Cerebellum | GM |
| 103 | 4.09 | -41 | 54 | 1 | L | Middle frontal gyrus | GM |
| 92 | 4.69 | -5 | 34 | -10 | L | Medial orbitofrontal cortex | GM |
| 88 | 4.26 | 30 | -42 | 49 | R | Superior parietal lobule | WM |
| 80 | 4.45 | 25 | 64 | 12 | R | Superior frontal gyrus | GM |
| 72 | 4.02 | 10 | -59 | 39 | R | Precuneus | GM |
| 71 | 3.72 | 52 | 13 | 24 | R | Inferior frontal gyrus | GM |
| 61 | 3.75 | -8 | -64 | 4 | L | Lingual gyrus | GM |
| 58 | 4.63 | 39 | -21 | 32 | R | Postcentral gyrus | WM |
| 55 | 4.43 | -25 | -97 | -4 | L | Cuneus | GM |

| | | | | | | | |
|---|---|---|---|---|---|---|---|
| 54 | 3.98 | -43 | 14 | -38 | L | Middle temporal gyrus | GM |
